# Accelerated brain aging towards transcriptional inversion in a zebrafish model of familial Alzheimer’s disease

**DOI:** 10.1101/262162

**Authors:** Nhi Hin, Morgan Newman, Jan Kaslin, Alon M. Douek, Amanda Lumsden, Xin-Fu Zhou, Alastair Ludington, David L. Adelson, Stephen Pederson, Michael Lardelli

## Abstract

Alzheimer’s disease (AD) develops silently over decades. We cannot easily access and analyse pre-symptomatic brains, so the earliest molecular changes that initiate AD remain unclear. Previously, we demonstrated that the genes mutated in early-onset, dominantly-inherited familial forms of AD (fAD) are evolving particularly rapidly in mice and rats. Fortunately, some non-mammalian vertebrates such as the zebrafish preserve fAD-relevant transcript isoforms of the *PRESENILIN* (*PSEN1* and *PSEN2*) genes that these rodents have lost. Zebrafish are powerful vertebrate genetic models for many human diseases, but no genetic model of fAD in zebrafish currently exists. We edited the zebrafish genome to model the unique, protein-truncating fAD mutation of human *PSEN2*, K115fs. Analysing the brain transcriptome and proteome of young (6-month-old) and aged, infertile (24-month-old) wild type and heterozygous fAD-like mutant female sibling zebrafish supports accelerated brain aging and increased glucocorticoid signalling in young fAD-like fish, leading to a transcriptional ‘inversion’ into glucocorticoid resistance and vast changes in biological pathways in aged, infertile fAD-like fish. Notably, one of these changes involving microglia-associated immune responses regulated by the ETS transcription factor family is preserved between our zebrafish fAD model and human early-onset AD. Importantly, these changes occur before obvious histopathology and likely in the absence of Aβ. Our results support the contributions of early metabolic and oxidative stresses to immune and stress responses favouring AD pathogenesis and highlight the value of our fAD-like zebrafish genetic model for elucidating early changes in the brain that promote AD pathogenesis. The success of our approach has important implications for future modelling of AD.

## Introduction

Alzheimer’s disease (AD) is the leading cause of dementia, a condition characterised by the progressive decline of memory and cognition. Like other neurodegenerative diseases, AD affects diverse cellular processes in the brain, including mitochondrial function^1^,^2^, metal ion homeostasis^3–5^, lipid metabolism^6–8^, immune responses^9^,^10^, synaptic transmission^11^, and protein folding and trafficking^12^,^13^. Dysregulation of these processes eventually results in severe atrophy of several brain regions (reviewed by Braak and Braak ^14^ and Masters et al. ^15^). Consequently, late stages of AD are likely to be much more difficult to treat than earlier stages of AD, contributing to our failure to discover ameliorative drugs^16^.

The pathological processes that result in AD are likely to initiate decades before clinical symptoms arise. Increased levels of soluble amyloid beta (Aβ) peptides in the cerebrospinal fluid and blood plasma is one of the earliest markers of both sporadic and familial forms of AD, preceding disease onset by 20-30 years^17^,^18^, while vascular changes are likely to occur even earlier^19^. Additionally, individuals possessing highly penetrant, dominant mutations in genes linked to the familial form of AD (fAD) such as *PSEN1* show structural and functional changes in their brains as early as 9 years of age, despite being cognitively normal^20^,^21^. Similar findings are evident in young adults carrying the ε4 allele of *APOE,* the major risk gene for the sporadic form of AD^22^. To prevent AD, we must identify the stresses underlying these early pathological changes. However, detailed molecular analysis of the brains of asymptomatic young adult fAD mutation carriers is currently impossible.

Analysing high-throughput ‘omics data (e.g. transcriptomic, proteomic) is a comprehensive and relatively unbiased approach for studying complex diseases like AD. Over the past decade, numerous post-mortem AD brains have been profiled using microarray and RNA-seq technologies, exposing an incredibly complex and interconnected network of cellular processes implicated in the disease^23^,^24^. Unfortunately, analysing post-mortem AD brains does not discern which cellular processes are responsible for initiating the cascade of events leading to AD.

Animal models can assist exploration of the early molecular changes that promote AD. However, early “knock-in” mouse models that attempted to model the genetic state of human fAD showed no obvious histopathology^25–27^. Modern ‘omics technologies provide molecular-level descriptions of disease states, but these technologies were not available when the early knock-in models were made. Subsequent transgenic models of AD constructed with multiple genes and/or mutations have displayed (what are assumed to be) AD-related histopathologies and these have also been analysed by ‘omics methods. However recent analysis of brain transcriptomes from five different transgenic AD models showed little concordance with human, late onset, sporadic AD brain transcriptomes. Worse still, none of the models were concordant with each other^28^.

Surprisingly, there has not yet been a detailed molecular investigation of the young adult brains of any animal model closely imitating the human fAD genetic state - i.e. heterozygous for a fAD-like mutation in a single, endogenous gene. Previously, we used zebrafish to analyse the unique, frameshifting fAD mutation of human *PRESENILIN2 (PSEN2),* K115fs, that inappropriately mimics expression of a hypoxia-induced truncated isoform of PSEN2 protein, PS2V^29–32^. Mice and rats have lost the ability to express PS2V^33^ (and the fAD genes of these rodents are evolving more rapidly than in many other mammals^33^), but in zebrafish, this isoform is expressed from the animal’s *psen1* gene^32^. Consequently, to model and explore early changes in the brain driving AD pathogenesis, we have now used gene-editing technology to introduce a K115fs-equivalent mutation into the zebrafish *psen1* gene, K97fs. In this paper, we analyse RNA-seq and mass spectrometry data collected from young adult (6-month-old) and aged, infertile (24-month-old) adult mutant and wild type zebrafish brains to comprehensively assess gene and protein expression changes in the brain due to aging and this fAD-like genetic state. At the molecular level, we find that the young fAD-model brains show elements of accelerated aging while aged fAD-like brains appear to ‘invert’ into a distinct and presumably pathological state. Our results emphasise the difficulty of understanding the early molecular progression of AD by examining overtly diseased brains and highlight the importance of accurate genetic models of fAD for elucidating mechanisms of AD pathogenesis.

## Results

Gene editing in zebrafish to produce the *psen1* K97fs mutation is described in the **Materials and Methods** section and in **Supplementary Methods 1**. To determine whether the K97fs mutation in the zebrafish *psen1* gene induces gene and protein expression changes, we removed whole-brains of *psen1*^K97fs/+^ (mutant) and *psen1*^+/+^ (wild type) adult zebrafish for total RNA sequencing (RNA-seq) and label-free tandem mass spectroscopy (LC-MS/MS) when zebrafish were 6 months (young) and 24 months (aged) old. We used three biological replicates to represent each of the four experimental conditions (young wild type, young mutant, aged wild type, aged mutant), and performed pairwise comparisons between experimental conditions to determine differentially expressed (DE) genes and differentially abundant (DA) proteins (**Figure 1**). Full lists of DE genes and DA proteins are provided in **Supplementary Table 1** and **Supplementary Table 2**.

**Figure 1.**
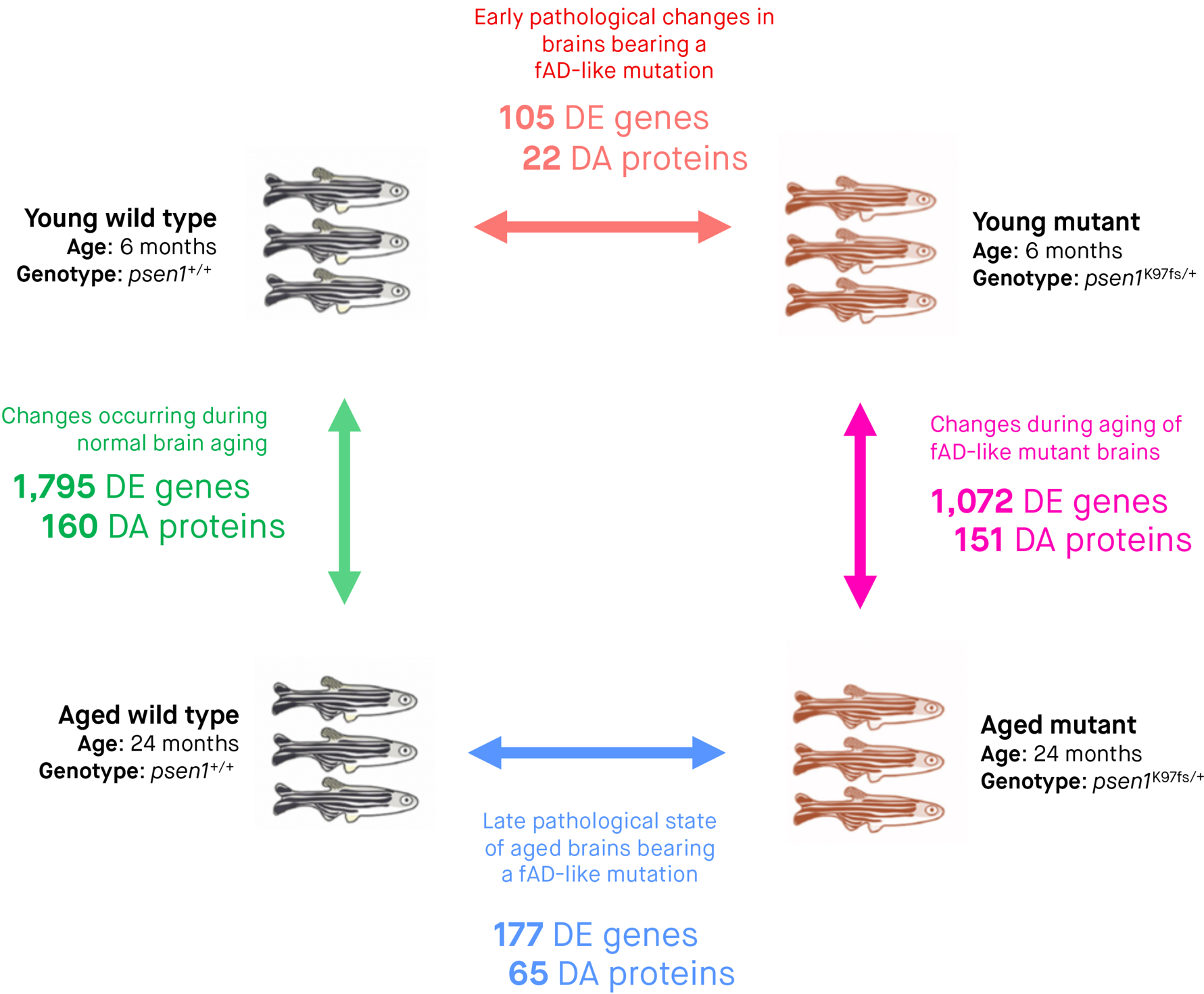
**Summary of experimental groups, comparisons, and differentiaMy expressed (DE) genes and differentially abundant (DA) proteins**. Three biological replicates (whole zebrafish brains) were subjected to RNA-seq and LC-MS/MS for each of the four experimental conditions. Arrows indícate pairwise comparisons (to identify DE genes and DA proteins) between experimental conditions. The numbers of DE genes and DA proteins determined from RNA-seq and LC-MS/MS analyses are indicated underneath the arrow for each comparison. We considered genes to be DE and proteins to be DA if the False Discovery Rate [FDR]-adjusted *p*-value of their moderated f-test *(limma)* was below 0.05. All zebrafish of the same age are siblings raised in the same tank.

### A familial AD-like mutation in zebrafish induces age-dependent gene expression changes

#### Early gene expression changes

The brains of children or young adults carrying fAD mutations display morphological and functional differences compared to age-matched individuals without these mutations^20^,^21^. Because of this, we hypothesised that gene expression in the brains of young adult (6-month-old) zebrafish carrying a fAD-like mutation would also be altered when compared to wild type zebrafish siblings. Overall, we find supporting evidence for 105 genes that are differentially expressed in young mutant brains relative to wild type brains (65 up-regulated, 40 down-regulated; FDR-adjusted *p*-value < 0.05) (**Supplementary Fig. 1**). Of these 105 genes, 65 have an log_2_ fold change greater than 0.5 or less than −0.5 in the ‘young mutant vs. young wild type’ comparison (**Figure 2A**). By examining the expression of these genes in the other three comparisons described in **Figure 1**, we observe two important phenomena:

1. **Accelerated aging genes are associated with increased immune response:** 62% (65/105) of the genes that are DE in 6-month-old mutant brains (‘young mutant vs. young wild type’) show the same direction of expression change during normal aging (‘aged wild type vs. young wild type’). However, far more genes are DE during normal aging (1,795 compared to 105). This suggests that the 6-month-old mutant brains may demonstrate accelerated aging for a subset of cellular functions. As an initial step to uncover these altered cellular functions, we applied functional enrichment analysis on these 65 genes and discovered significant enrichment in an MSigDB gene set relating to immune response genes that are up-regulated following lipopolysaccharide treatment “GSE9988 LPS VS VEHICLE TREATED MONOCYTE UP” (Bonferroni adjusted *p*-value 0.000948) (**Supplementary Table 3**).
2. **Age-dependent ‘inversión’ pattern:** A subset of 63 genes with increased expression in 6-month-old mutant brains (‘young mutant vs. young wild type’) show decreased expression in 24-month-old mutant brains (‘aged mutant vs. aged wild type’). We call this expression pattern an age-dependent ‘inversion’ between mutant and wild type brains. We explore the biological relevance of the genes involved in this inversion pattern in the transcriptome later.

**Figure 2.**
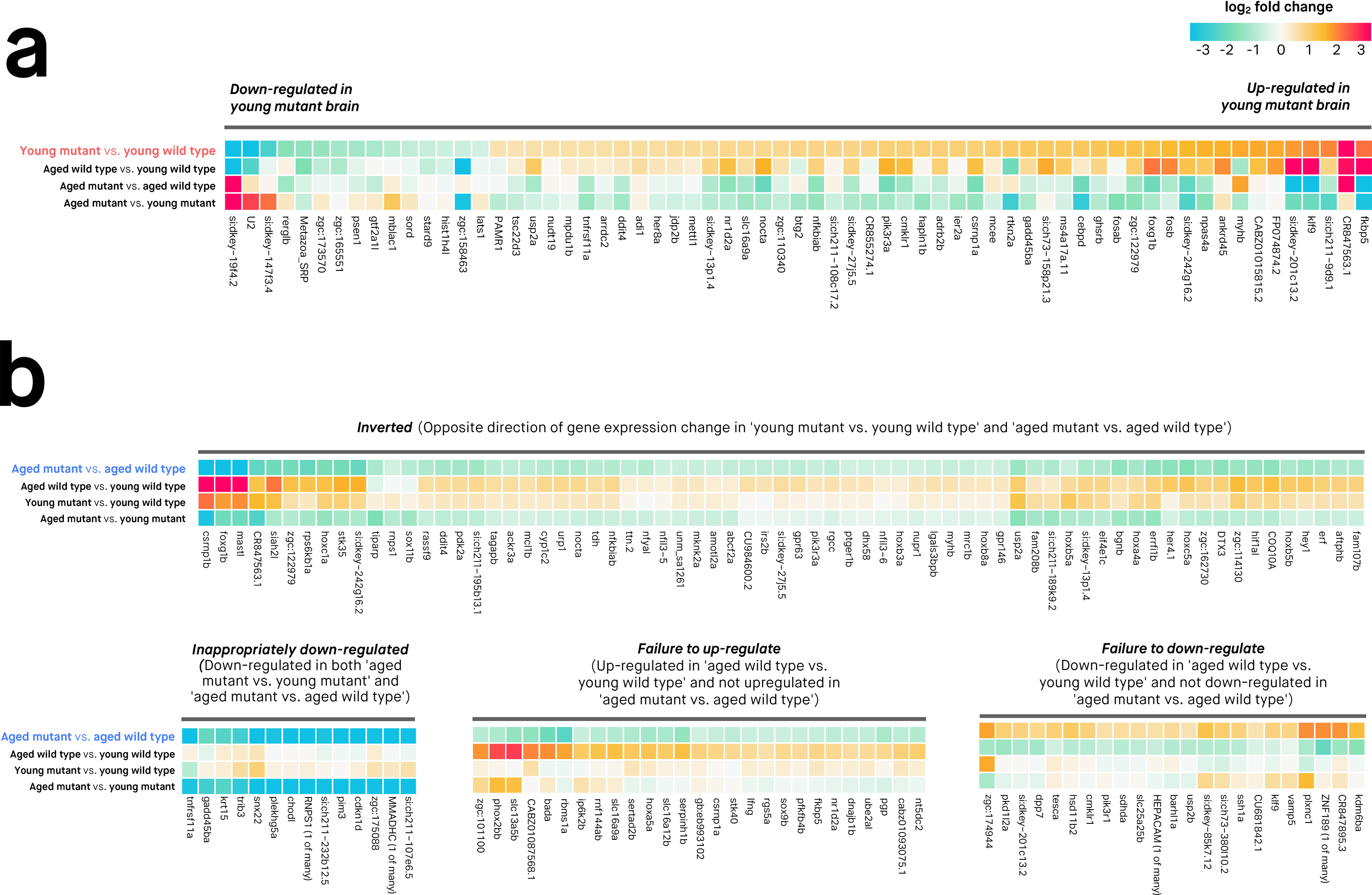
**Gene expression changes in the brains of fAD-like mutant (*psen1*^K97fs/+^) zebrafish compared to wild type (*psen1*^+/+^) siblings.** Genes shown in the heatmaps only include those with FDR-adjusted *p*-value < 0.05 and absolute log_2_ fold change > 0.5. A. Differentially expressed genes between young (6-month-old) mutant and wild type zebrafish brains. B. Differentially expressed genes between infertile, aged (24-month-old) mutant and wild type zebrafish brains. The differentially expressed genes are separated into clusters based on gene expression changes across the four comparisons. Overall, note the similar expression changes in 'young mutant vs. young wild type' and 'aged wild type vs. young wild type' and the contrast of these to comparisons involving aged mutants. This illustrates the accelerated brain aging in young mutant brains and the "inverted" gene expression pattern of aged mutant brains.

#### Late gene expression changes

By comparing gene expression in 24-month-old mutant and wild type zebrafish brains, we can gain insight into a putatively pathological transcriptomic state present in the brains of aged zebrafish carrying a fAD-like mutation. We find supporting evidence for 177 genes that are differentially expressed in mutant brains relative to wild type brains (139 down-regulated, 38 up-regulated, FDR-adjusted *p*-value< 0.05) (**Figure 2B**; **Supplementary Fig. 1**). Note that not all of these genes are shown in **Figure 2B**, which only includes genes with log_2_ fold change values greater than 0.5 or less than −0.5. To allow for easier interpretation of these 177 genes, we used hierarchical clustering to separate them into groups with distinct expression patterns based on all four brain-types:

▪ **Inverted (63 genes)**: Defined as genes showing opposite fold-changes in young mutant brains (‘young mutant vs. young wild type’) compared to aged mutant brains (‘aged mutant vs. aged wild type’). To be included in this group, genes were required to have an FDR-adjusted *p*-value < 0.05 in either the ‘young mutant vs. young wild type’ or ‘aged mutant vs. aged wild type’ comparison and an unadjusted *p*-value < 0.05 in the other comparison.
▪ **Inappropriately down-regulated (57 genes)**: Defined as genes that are down-regulated in the ‘aged mutant vs. young mutant’ and ‘aged mutant vs. aged wild type’ comparisons.(FDR-adjusted *p*-value 0.05 in both).
▪ **Failure to up-regulate (94 genes)**: Defined as genes that are up-regulated during normal aging (FDR-adjusted *p*-value < 0.05 in the ‘aged wild type vs. young wild type’ comparison) but not up-regulated in the ‘aged mutant vs. aged wild type’ comparison.
▪ **Failure to down-regulate (26 genes)**: Defined as genes that are down-regulated during normal aging (FDR-adjusted *p*-value < 0.05 in the ‘aged wild type vs. young wild type’ comparison) but not down-regulated in the ‘aged mutant vs. aged wild type’ comparison.

To determine whether these different component groups of the gene expression patterns are biologically relevant, we assessed each group’s functional enrichment using Gene Ontology terms, MSigDB gene sets, and Reactome and Interpro pathways (summarised in **Supplementary Table 3**; full results in **Supplementary Table 4**). Overall, we find statistically significant enrichment (Bonferroni adjusted *p*-value < 0.05) for all groups except for the ‘failure to down-regulate group. The ‘inverted’ group is significantly enriched in several gene sets related to stress and immune response; the ‘inappropriately down-regulated’ group is significantly enriched in developmental transcription factors including homeobox genes; the ‘failure to up-regulate’ group is significantly enriched in immune response.

#### Gene expression changes due to brain aging

By comparing gene expression in 24-month-old and 6-month-old brains, it is possible to identify genes that show altered expression during normal aging and the aberrant molecular aging of the fAD-like mutant fish. 1,795 genes show altered expression levels with normal brain aging in zebrafish (‘aged wild type vs. young wild type’), while 1,072 genes show altered expression in mutant brains as they age (‘aged mutant vs. young mutant’) (FDR-adjusted *p*-value < 0.05). 525 genes overlap in these two sets, and can be considered an ‘aging signature’ showing statistically significant fold changes in the same direction during both wild type and mutant brain aging. These genes are enriched in gene ontology terms related to immune function (**Supplementary Table 3**) indicating that some immune responses that are associated with normal brain aging are still preserved in mutant zebrafish.

### The gene expression changes observed are likely not due to changes in the proportions of brain cell types

It is possible that changes in the proportions of different cell types in the brain could result in genes being falsely interpreted as differentially expressed. To ensure that our observations of differential gene expression are not an artefact of changes in the proportions of the major brain cell types (e.g. astrocytes, microglia, neurons, oligodendrocytes), we checked that the average expression for sets of marker genes representing each of the major brain cell types was approximately constant across the samples in each biological condition (young wild type, young mutant, aged wild type, aged mutant). Representative marker genes for microglia were obtained from Oosterhof et al.^34^ while gene markers for astrocytes, neurons, and oligodendrocytes were obtained from Lein et al.^35^ The number of genes used to calculate the average gene expression (in logCPM) was 41 (astrocyte), 99 (microglia), 77 (neuron) and 78 (oligodendrocyte). Overall, the average expression of gene markers for the major neural cell types is approximately constant in each of the biological conditions, and no obvious outlier samples are evident (**Supplementary Fig. 3**).

### Regulation of gene expression changes in fAD-like mutant zebrafish brains differs from normal brain aging

A transcription factor can regulate (activate or repress) gene expression by binding to a specific DNA motif in the promoter region of a gene. We hypothesised that changes in gene expression during normal aging or differences in gene expression between mutant and wild type brains would be driven by differences in transcription factor activity. To test this, we examined gene promoter regions for enriched motifs corresponding to known transcription factor binding sites (summarised in **Supplementary Table 5**; full results in **Supplementary Table 6**). Overall, we find:

1. **Numerous known transcription factors likely drive the gene expression changes that occur during normal zebrafish brain aging**. As wild type brains age, the genes which are differentially expressed are significantly enriched in many known motifs. These motifs correspond to binding sites for interferon regulatory factors (e.g. IRF1, IRF2, IRF8); a binding site for the the PU. 1-IRF8 complex; an interferon-stimulated response element (ISRE); and binding sites for various transcription factors important for essential cellular processes like proliferation, differentiation, and apoptosis (Atf3, Fra2, Ets-distal, AP-1, Fra1, JunB, BATF, and ZNF264). This supports the idea that numerous transcription factors contribute to the coordinated gene expression changes observed during aging.
2. **Altered glucocorticoid signalling in mutant zebrafish brains is likely to contribute to brain pathology in fAD-like mutants**. Promoters of genes that are differentially expressed in the ‘aged mutant vs. aged wild type’ comparison are significantly enriched in the glucocorticoid receptor element motif (GRE) (Bonferroni *p*-value = 0.0057). Interestingly, the subset of genes showing inappropriate downregulation (down-regulated in the ‘aged mutant vs. young mutant’ and ‘aged mutant vs. aged wild type’ comparisons) is even more enriched in the GRE motif (Bonferroni *p*-value = 0.0001), suggesting that genes that are normally activated by glucocorticoid signalling during aging may not be activated in aged mutant brains. This altered glucocorticoid signalling appears to be present even in young zebrafish brains, as genes showing inverted behaviour (opposite direction of differential expression in ‘young mutant vs. young wild type’ and ‘aged mutant vs. aged wild type’ comparisons) are also enriched in the GRE motif (Bonferroni *p*-value = 0.0047). Because these inverted genes tend to show high expression in young mutant brains (i.e. up-regulated in the ‘young mutant vs. young wild type’ comparison) and low expression in aged mutant brains (i.e. down-regulated in the ‘aged mutant vs. aged wild type’ comparison), this suggests that young mutant zebrafish brains may initially exhibit abnormally increased glucocorticoid signalling, while aged mutant brains later exhibit abnormally decreased glucocorticoid signalling. Notably, the inverted genes containing a GRE motif in their promoters include *COQ10A* (encodes Coenzyme Q10, a key component of the electron transport chain and free-radical scavenging antioxidant); *pik3r3a* (encodes regulatory subunit gamma of phosphoinositide 3- kinase, an enzyme that interacts with insulin growth factor 1 receptor among other proteins); *mmadhc* (encodes a protein involved in an early and essential step of vitamin B12 metabolism), *plk3* (polo-like kinase 3, involved in stress response and double-stranded DNA repair), and *fkbp5* (encodes FK506 binding protein, involved in regulating immune and stress responses, protein trafficking and folding, and glucocorticoid receptor regulation). A list of zebrafish genes containing the GRE promoter motif is provided in **Supplementary Table 7**.

### expression changes in the fAD-like mutant indicate vast changes to cellular processes and pathways

A gene set is a group of genes that contribute to a predefined biological function, pathway, or state. A gene set test is an analysis used to evaluate whether a particular gene set is differentially expressed for a particular comparison. We used FRY to test whether ‘Hallmark’ gene sets from the Molecular Signatures Database (MSigDB)^36^ were differentially expressed in each of the four comparisons (**Figure 3, Supplementary Table 8**). Using an FDR-adjusted *p*-value < 0.05 to define a gene set as differentially expressed, we find:

1. **50 gene sets are differentially expressed during normal brain aging (‘aged wild type vs. young wild type’)** (middle row of heatmap, Figure 3A). This supports that many biological functions and pathways are altered during normal aging. For some gene sets, the proportion of genes that are up-regulated and down-regulated is similar (e.g. interferon alpha response, E2F targets, early estrogen response). However, other gene sets contain a predominance of up-regulated genes (e.g. epithelial mesenchymal transition, TNFA signalling via NFKB) or down-regulated genes (e.g. coagulation, reactive oxygen species pathway).
2. **22 gene sets are differentially expressed in young mutant brains ('young mutant vs. young wild type')** (top row of heatmap, **Figure 3A**). These 22 gene sets may represent earlier functional changes in the brain that occur due to a fAD-like mutation. The gene sets implicate diverse processes including Wnt/β-catenin signalling, early estrogen response, DNA repair, hedgehog signalling and fatty acid metabolism. Similar to the pattern of accelerated aging observed in **Figure 2**, we also observe that most of the gene sets up-regulated in young mutant brains are regulated in the same direction during normal aging. This is consistent with the idea that the biological changes in young mutant brains may partially recapitulate those that occur during normal brain aging.
3. **44 gene sets are differentially expressed between aged mutant and wild type brains** (bottom row, **Figure 3A**). These differentially expressed gene sets may represent the pathological state of aged zebrafish brains bearing a fAD-like mutation. Importantly, 21 of the 22 gene sets that were differentially expressed in young mutant brains ('young mutant vs. young wild type') remain altered also when these are aged ('aged mutant vs. aged wild type'). However, the proportions of up- and down-regulated genes tend to differ; notably, several gene sets containing a predominance of up-regulated genes in the young mutant brains contain a predominance of down-regulated genes in the old mutant brains. These ‘inverted’ gene sets include biological functions and pathways as diverse as Wnt/β-catenin signalling, early estrogen response, hedgehog signalling, androgen response, epithelial mesenchymal transition, DNA repair, apical surface, and TGF-β signalling.
4. **Aging in mutant brains is similar but distinct from aging in wild type brains.** The 50 gene sets differentially expressed during normal brain aging are also differentially expressed during mutant brain aging ('aged mutant vs. young mutant') (**Figure 3B**). However, proportions of up- and down-regulated genes differ from those in normal brain aging. This suggests that zebrafish brains bearing a fAD-like mutation may not properly regulate certain gene sets during aging (e.g. cholesterol homeostasis, adipogenesis, DNA repair, hypoxia, Wnt/β-catenin signalling).

**Figure 3.**
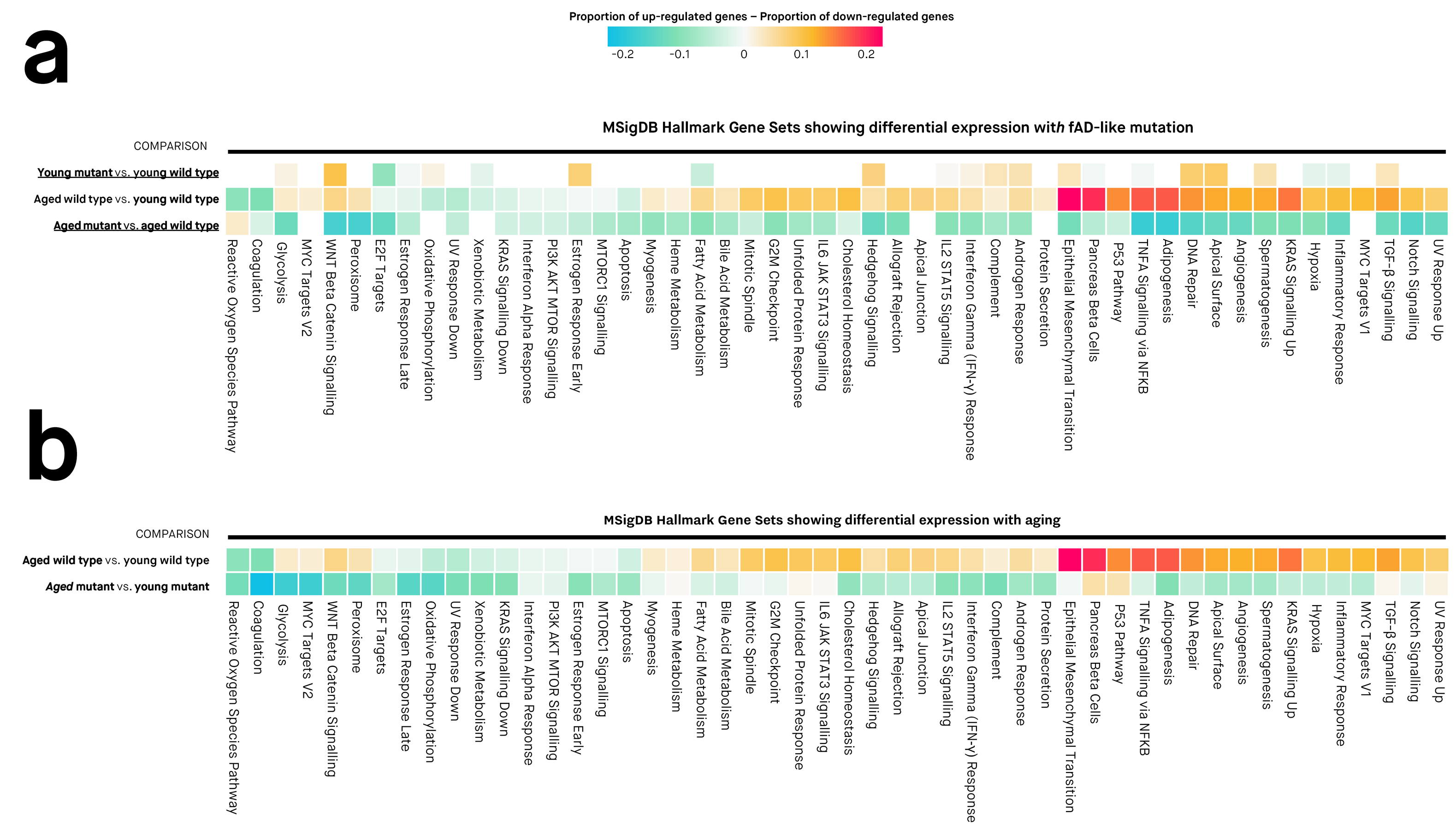
**Gene sets showing differential expression with aging and/or familial Alzheimer’s disease (fAD)-like mutation.** Colours of cells are proportional to the difference between the proportion of up- and down-regulated genes in a gene set. Differentially expressed gene sets have Mixed FDR below 0.05, indicating the gene set is likely to show statistically significantly altered (up or down) expression for a particular comparison. The genes in each gene set are defined using the "Hallmark" gene set collection at the Molecular Signatures Database (MSigDB).

### Altered protein abundances in fAD-like mutant zebrafish

Mutant zebrafish brains exhibit numerous transcriptional alterations linked to diverse cellular processes. However, since different mechanisms regulate the stability of RNA transcripts and proteins, it can be difficult to predict changes in protein abundance from RNA-seq data. We used LC-MS/MS to explore protein abundance changes in mutant zebrafish brains. Unfortunately, the resulting proteomic data are not directly comparable to the RNA-seq data due to only reliably quantifying 323 proteins from LC-MS/MS compared to 18,296 genes from RNA-seq (see **Supplementary Methods 2** and **Supplementary Fig. 4 and 5** for correlation analysis between the RNA-seq and LC-MS/MS data sets). Nevertheless, the proteomic data reveal that numerous proteins are differentially abundant across each of the four comparisons. Here, we focus on the comparisons between mutant and wild type brains in young and aged zebrafish brains (**Figure 4**, **Supplementary Fig. 2**).

**Figure 4.**
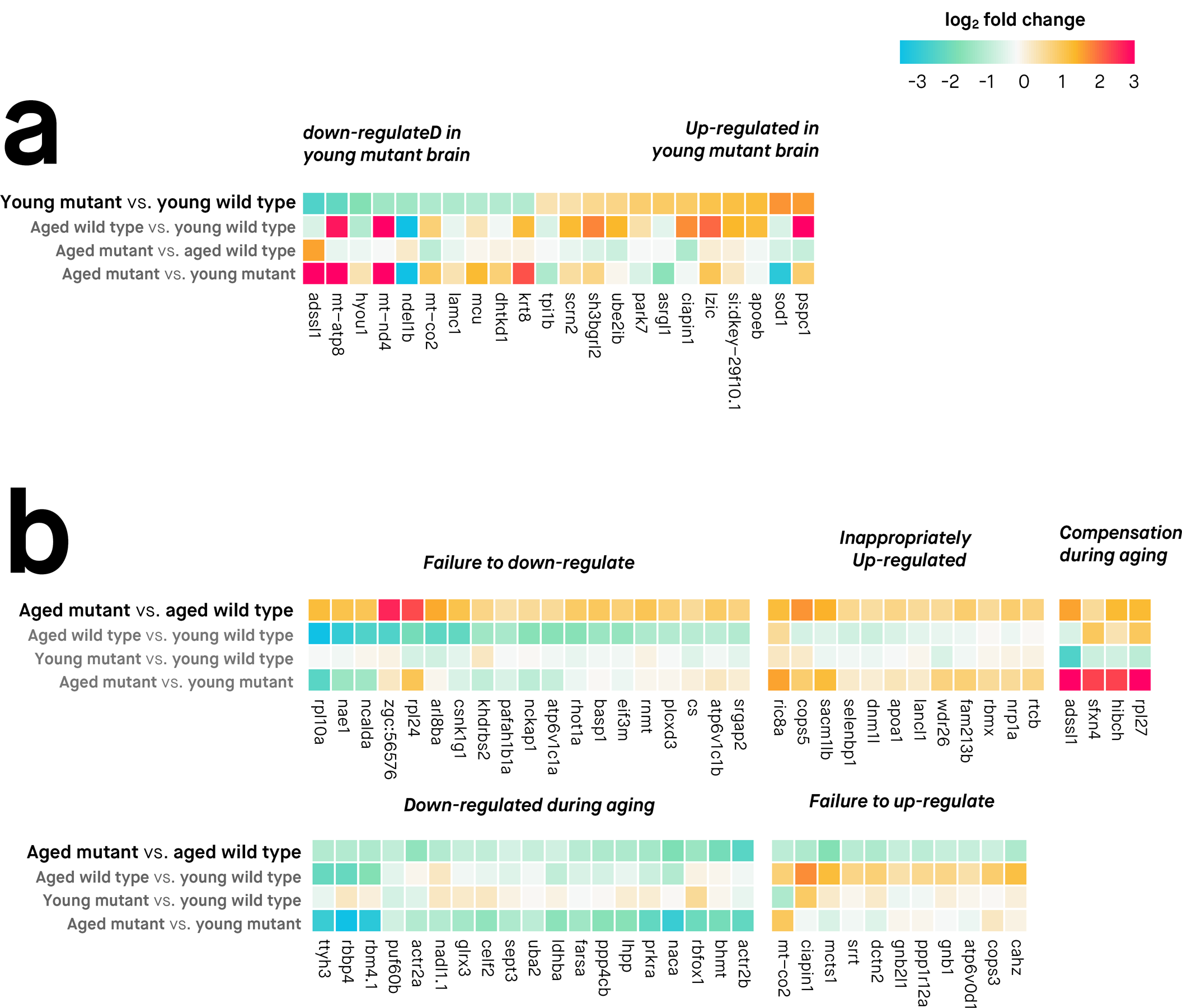
**Protein abundance changes in the brains of fAD-like mutant zebrafish compared to wild type siblings at 6 months (young) and 24 months (aged) of age.**Protein abundance was quantified at the peptide-level with LC-MS/MS (liquid chromatography tandem mass spectrometry) and differential abundance was assessed using moderated f-tests *(limma).* Differentially abundant proteins are defined as those with FDR-adjusted *p*-value < 0.05. Protein names were used to retrieve equivalent gene symbols for display purposes on these heatmaps. **A. Differentially abundant proteins between young mutant and wild type zebrafish brains. B. Differentially abundant proteins between aged mutant and wild type zebrafish brains.** The proteins have been clustered according to their abundance changes across the four comparisons.

#### Early protein abundance changes

1. When zebrafish are 6 months old, 22 of the 323 detected proteins are differentially abundant between mutant and wild type brains ('young mutant vs. young wild type' comparison, **Figure 4A**) (12 up-regulated, 10 down-regulated, FDR-adjusted *p*-value < 0.05).
2. Remarkably, three of the 12 up-regulated proteins have well-established roles in neurodegenerative diseases: apolipoprotein Eb (encoded by the zebrafish *apoeb* gene, orthologous to the major human genetic risk gene for sporadic AD, *APOE);* superoxide dismutase (encoded by the zebrafish *sod1* gene, orthologous to the human *SOD1* gene mutated in familial amyotrophic lateral sclerosis); and protein DJ-1 (encoded by the zebrafish *park7* gene, orthologous to the human *PARK7* gene mutated in familial Parkinson's disease).
3. Several mitochondrial proteins (cytochrome c oxidase subunit 2, NADH-ubiquinone oxidoreductase chain 4, ATP synthase protein 8, mitochondrial calcium uniporter protein, probable 2-oxoglutarate dehydrogenase E1) are decreased in 6-month-old mutant brains. This is consistent with the alterations in energy metabolism and oxidative stress previously identified as early events in AD pathogenesis^37^,^38^.

#### Late protein abundance changes

1. When zebrafish are 24 months of age, 65 of the 323 proteins are differentially abundant between mutant and wild type brains ('aged mutant vs. aged wild type' comparison, **Figure 4B**) (35 up-regulated, 30 down-regulated FDR-adjusted *p*-value < 0.05).
2. As with the differentially expressed genes in Figure 2, the differentially abundant proteins for this comparison are also grouped according to their abundance changes in the four comparisons. The 'failure to down-regulate' and 'failure to up-regulate' clusters represent proteins that normally decrease or increase during normal brain aging, but fail to do so during the aberrant aging of mutant brains. The 'inappropriately up-regulated' and 'compensation during aging' clusters represent proteins that tend to not be differentially abundant at 6-months (except for *adssl1),* but increase during aging of mutant brains, finally becoming differentially abundant in the aged mutant brains.
3. In contrast to the differentially expressed genes in Figure 2, many of the protein abundance changes appear to be specific to a particular age group (i.e. either 6-month-old or 24-month-old). Only three proteins (encoded by the *adssl, mt-co2, ciapin1* genes) are differentially abundant at both 6 months and 24 months of age. Interestingly, several proteins that are differentially abundant at 6 months (including apolipoprotein Eb, superoxide dismutase and protein DJ-1) are no longer differentially abundant at 24 months.

Overall, the protein abundance results provide a complementary perspective to the differentially expressed genes and gene sets identified earlier.

### An interconnected view of altered gene expression in fAD mutant zebrafish brains

Our results support the idea that aged mutant zebrafish brains contain dysregulated genes that are involved in diverse cellular processes, raising several key questions.
1. What particular cellular processes are critical for the pathological state observed in aged fAD-like mutant brains?
2. Are these pathological changes similar to those in human brains with fAD?

To address these questions, we compared gene expression patterns in zebrafish and human brains by constructing co-expression networks of genes. The zebrafish co-expression network was constructed using the RNA-seq data described earlier; the human co-expression network was constructed using a microarray-based dataset from Antonell et al. ^24^ (GEO accession number GSE39420). The Antonell et al. dataset includes patients with fAD caused by *PSEN1* mutations and patients with early-onset AD lacking *PSEN1* mutations. This allows us to compare co-expression patterns in our mutant zebrafish to*PSEN1*-linked AD and the more general cases of early-onset AD.

#### Constructing gene co-expression networks in zebrafish and human

When constructing the co-expression networks, we only included genes that were orthologs in humans and zebrafish. Whilst there are many methods for constructing a co-expression network of gene expression^39^, we used the weighted gene co-expression network analysis (WGCNA) method^40^. WGCNA has previously been used to group genes expressed in the brain into “modules” that are associated with biological functions^41–45^. The zebrafish brain co-expression network is shown in **Figure 5A**, and the human brain co-expression network is provided in **Supplementary Fig. 6**.

**Figure 5.**
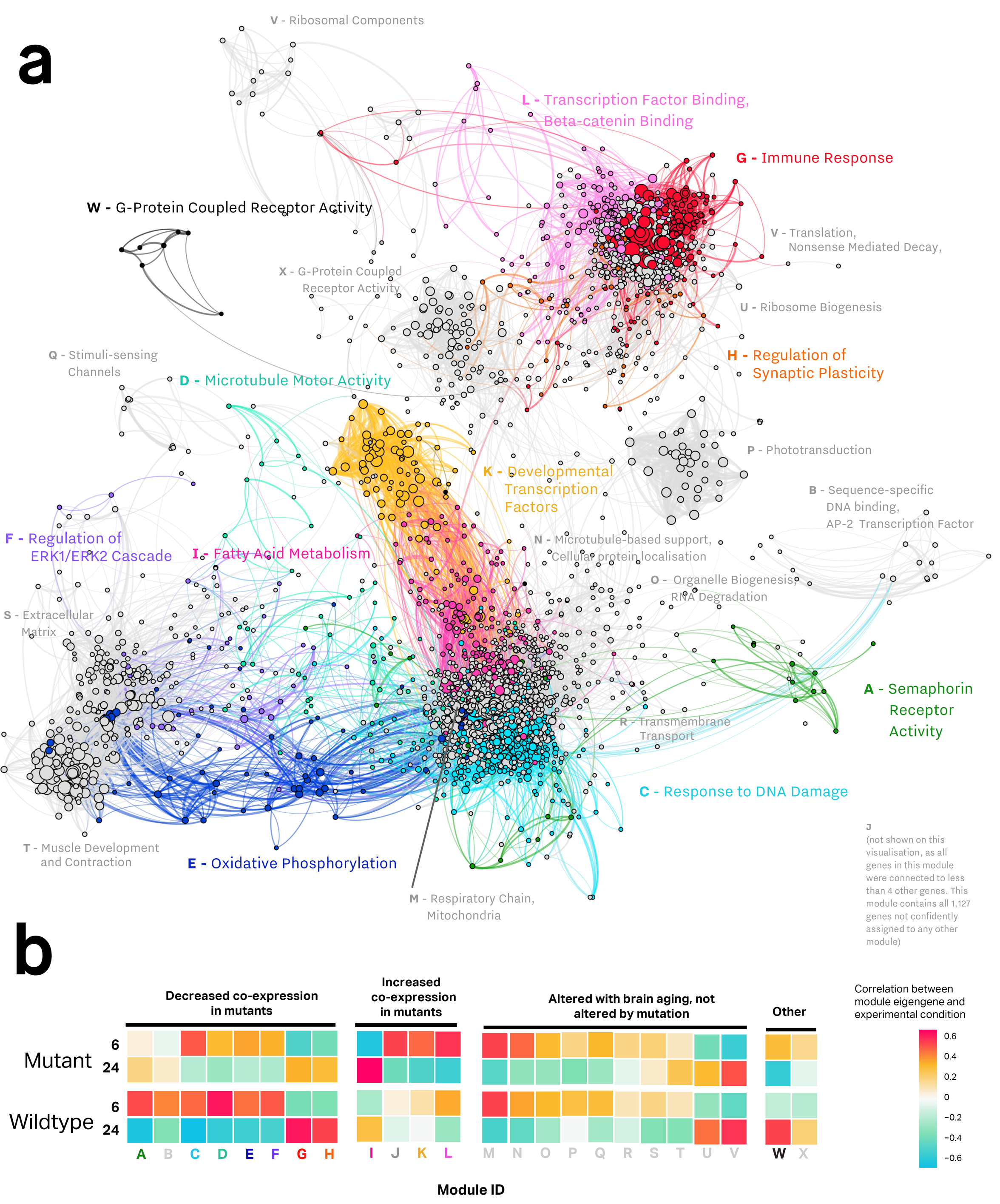
**Zebrafish brain gene co-expression network. A. Gene co-expression network visualisation.** Each node represents one gene, with node size proportional to the number of connected nodes (co-expressed genes). Edges represent co-expression between two genes, with edge weight proportional to the strength of co-expression. The co-expression network is a signed adjacency matrix constructed from gene expression data from wild type and mutant zebrafish brains at 6 and 24 months of age. Only nodes with at least four connections are shown. Alphabet letters correspond to "modules" of co-expressed genes. Modules that show altered gene co-expression patterns in mutant zebrafish brains relative to wild type zebrafish brains are coloured. **B. Gene expression patterns of modules in the gene co-expression network across mutant and wild type zebrafish brains at 6 months and 24 months of age.** Colours of cells are indicate the hybrid Pearson-robust correlations between the overall gene expression in a module (summarised using the first principal component) and experimental condition encoded as a binary variable (6-month-old mutant, 24-month-old mutant, 6-month-old wild type, 24-month-old wild type). Modules showing potentially altered expression patterns during mutant aging compared to wild type aging are labelled with coloured text, with colours corresponding to module colours in **A.**

#### Identification of biologically relevant modules in the co-expression networks

We identified 23 modules (i.e. groupings of genes) in the zebrafish brain co-expression network containing between 79 and 818 genes each, and 13 modules in the human brain co-expression network containing between 62 and 921 genes each. We used two methods to confirm that most modules represented functional relationships between genes: enrichment analysis (for identifying enriched biological functions and enriched promoter motifs), and correlating modules with particular traits of interest (age and/or*psen1* genotype).

#### Correlating modules with zebrafish traits

By correlating modules with particular zebrafish traits (age and *psen1* genotype), we identified the modules showing evidence of altered expression patterns in mutant zebrafish brains. Of the 23 modules identified here in the zebrafish brain (as **A** to **X**), 13 show evidence of disruption in the mutant brains compared to wild type brains (**Figure 5B**). Eight modules (from **A** to **H**) show decreased co-expression in mutant zebrafish brains, while six modules (from **I** to **L**) show increased co-expression in mutant zebrafish brains.

#### Enrichment Analysis

A summary of the biological relevance of each module in the zebrafish co-expression network is provided in **Table 1** (full enrichment analysis results in **Supplementary Tables 9, 10 and 11**). Overall, zebrafish and human network modules show significant enrichment in functional categories (e.g. Gene ontology terms, MSigDB gene sets, with Bonferroni-adjusted *p*-value < 0.05), supporting the idea that these modules are likely to represent biologically relevant groupings of genes. Some of the biological functions represented by different modules in the zebrafish brain include: immune response (represented by module **G**), oxidative phosphorylation (represented by module **E**), translation and ribosomal components (represented by module **V**), G-protein coupled receptor activity (represented by module **X**), and extracellular matrix (represented by module **S**). Only one module (**G**) was significantly enriched in promoter motifs (Bonferroni-adjusted *p*-values for enriched motifs < 0.05). The enriched promoter motifs in module G include several motifs recognised by the ETS transcription factor family (SpiB, ELF3 and ELF5; Bonferroni-adjusted *p*-values of 0.00424, 0.0106, and 0.0127 respectively) and interferon regulatory factor motifs (IRF3, IRF8, ISRE; Bonferroni-adjusted *p*-values of 0.0118, 0.0144, and 0.0260 respectively) (**Supplementary Table 12**).

**Table 1.** **Summary of modules in a co-expression gene expression network constructed from zebrafish RNA-seq data and their preservation in an independent human brain microarray data set.** The Ζ-Summary preservation score is a statistic that aggregates various *Ζ*-statistics obtained from permutation tests of the co-expression network to test whether network properties such as density and connectivity in the zebrafish co-expression network are preserved in an independent co-expression network constructed from human brain gene expression data^24^. In this analysis, 200 permutations were used. *Ζ*-summary scores less than 2 indicate no preservation, while scores between 2 and 10 indicate weak-to-moderate evidence of preservation. The top functional enrichment and cell type marker enrichment terms are used to give insight into possible biological functions represented within each module. Cell type marker enrichment gene sets are from MSigDB, while functional enrichment terms are from Gene Ontology and MSigDB gene sets. The “Random” module is a random sample of 1,000 genes in the zebrafish co-expression network expected to show non-significant preservation (*Ζ*-summary < 2) in the human co-expression network.

| Module | Number of Genes | Z-Summary Preservation Score | Top Functional Enrichment Terms (FDR $p$-value < 0.05) | Cell Type Marker Enrichment (FDR $p$-value < 0.05) |
| --- | --- | --- | --- | --- |
| **A** | 89 | 1.42 | – | – |
| **B** | 106 | -0.31 | Wnt Signaling Pathway, AP-2 Transcription Factor Family | – |
| **C** | 440 | -0.57 | Zinc Finger C2H2 | – |
| **D** | 273 | 2.16 | – | – |
| **E** | 132 | 0.53 | Oxidative Phosphorylation, Fatty Acid Beta-Oxidation | Astrocyte |
| **F** | 89 | 0.53 | Transmembrane Receptor Protein Tyrosine Kinase Activity, Regulation of ERK1 and ERK2 Cascade | – |
| **G** | 334 | 3.83 | Immune Response | Microglia |
| **H** | 79 | 3.72 | Regulation of Synaptic Plasticity, Synaptic Signaling | CA1 Pyramidal Neuron |
| **I** | 306 | 0.59 | – | – |
| **J** | 1178 | 4.91 | – | – |
| **K** | 126 | 0.37 | Glycinergic Synaptic Transmission, Developmental Transcription Factors bound by Suz12 | – |
| **L** | 267 | 0.98 | Circadian Clock, β-Catenin Binding | – |
| **M** | 714 | -0.23 | Mitochondrial Respiratory Chain | – |
| **N** | 243 | 0.22 | – | – |
| **O** | 532 | -0.99 | DNA Repair, Cytoplasmic Part | – |
| **P** | 85 | -0.68 | Visual Perception, Phototransduction | – |
| **Q** | 126 | 1.33 | – | – |
| **R** | 149 | 0.85 | Glycosylphosphatidylinositol (GPI)-anchor biosynthesis | – |
| **S** | 217 | 2.22 | Extracellular Matrix | – |
| **T** | 220 | 1.16 | ECM-Receptor Interaction, Muscle Structure Development | – |
| **U** | 117 | 0.27 | – | – |
| **V** | 818 | 0.38 | Translation, Ribosome, Nonsense Mediated Decay | – |
| **W** | 136 | 1.47 | – | – |
| **X** | 342 | 6.86 | G-Protein Coupled Receptor Activity, Synaptic Signaling | CA1 Pyramidal Neuron |
| **Random** | 1000 | 0.64 | – | – |

#### Several pathological changes in mutant zebrafish brains are similar to those in human fAD brains

There are several methods for assessing whether modules are preserved across two independent gene co-expression networks constructed using the same genes^46^. The most easily interpretable method is to compare directly the assignment of equivalent genes to modules identified in each network. The resulting overlap in gene co-expression patterns across the two networks can be visualised using a Sankey diagram (**Figure 6**). Overall, the gene co-expression patterns in the zebrafish brain appear to be broadly similar to the gene co-expression patterns in the human brain. A more sophisticated method of assessing module preservation involves using permutation-based *Ζ*-statistics to test whether certain properties of modules (e.g. density, connectivity) defined in one co-expression network are preserved in another network^46^. *Ζ*-statistics for each module property can be summarised into a *Ζ*-summary score, with *Ζ*-summary scores less than 2 indicating no module preservation, scores between 2 and 10 indicating weak to moderate module preservation, and scores above 10 indicating strong preservation^46^. When comparing zebrafish and human brain co-expression networks, six of the 23 zebrafish modules have *Ζ*-summary scores between 2 and 10, indicating weak to moderate preservation in the human co-expression network (**Table 1**, **Supplementary Table 13**). Importantly, three of these modules (**D, G, H**) also display altered co-expression in the mutant zebrafish brains (**Figure 5**), suggesting that at least several biological processes (“microtubule motor activity”, “immune response”, “regulation of synaptic plasticity”) altered in fAD-like mutant zebrafish brains may also be altered in the brains of humans with fAD. Notably, the module enriched in immune response functions, module **G** (*Ζ*-summary score 3.83), is also significantly enriched in ETS and IRF motifs.

**Figure 6.**
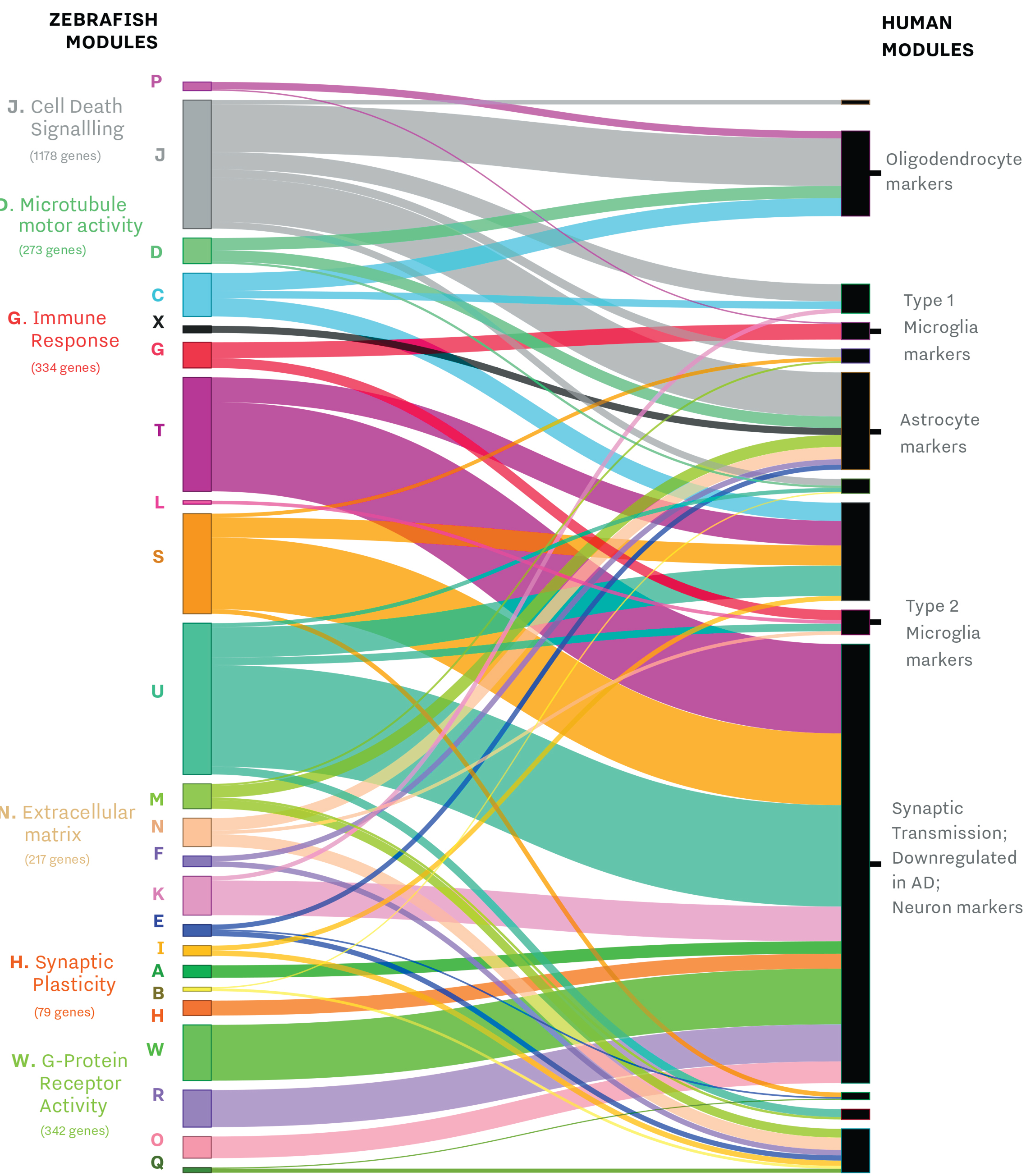
**Module overlap between co-expression networks constructed using zebrafish and human brain gene expression data.** Zebrafish and human co-expression networks were constructed using 7,118 genes that were orthologs in zebrafish and humans and expressed in brain gene expression data. Modules of co-expressed genes were separately identified for both the zebrafish and human co-expression networks, resulting in 23 modules in the zebrafish network (left) and 13 modules in the human network (right). Labelled zebrafish modules (J, D, G, N, H, W) have Z-summary preservation score > 2, indicating statistically significant weak-to-moderate preservation of these modules (i.e. genes in these modules still tend to be co-expressed) in the human brain co-expression network. Labels on zebrafish and network modules are based on the top functional enrichment terms found using Gene Ontology and MSigDB gene sets. See **Table 1** for more details on the Z-summary preservation scores and functional enrichment for each module in the zebrafish co-expression network.

#### In zebrafish, the molecular changes in aged mutant brains occur without obvious histopathology

Teleosts (bony fish) such as the zebrafish show impressive regenerative ability following tissue damage that includes repair of nervous tissue. Previous attempts to model neurodegenerative diseases in adult zebrafish have failed to show cellular phenotypes^47^. Also, zebrafish are thought unlikely to produce the Aβ peptide^48^ that many regard as central to AD pathological mechanisms^49^. The analyses described in this paper support that fAD mutations in the *PSEN* genes accelerate aspects of brain aging and promote a shift in aged mutant brains towards an altered, pathological state of gene and protein expression. We therefore made histopathological comparisons of aged (24 months) wild type and mutant brains equivalent to those used in our ‘omics analyses. Analysis of various brain regions using markers of aging, senescence and amyloid accumulation (lipofuscin, senescence-associated β-galactosidase, and congo red staining respectively) revealed no discernible differences (see **Supplementary Methods 3** and **Supplementary Fig. 7, 8** and **9**). This is consistent with the lack of neurodegenerative histopathology observed in a heterozygous knock-in model of a *PSEN* fAD mutation in mice^25^.

## Discussion

Using zebrafish with heterozygous fAD-like mutations in single endogenous genes appears to be useful for studying fAD pathogenesis at the molecular level. **Figure 7** summarises the main molecular changes that occur with aging and the fAD-like mutation.

**Figure 7.**
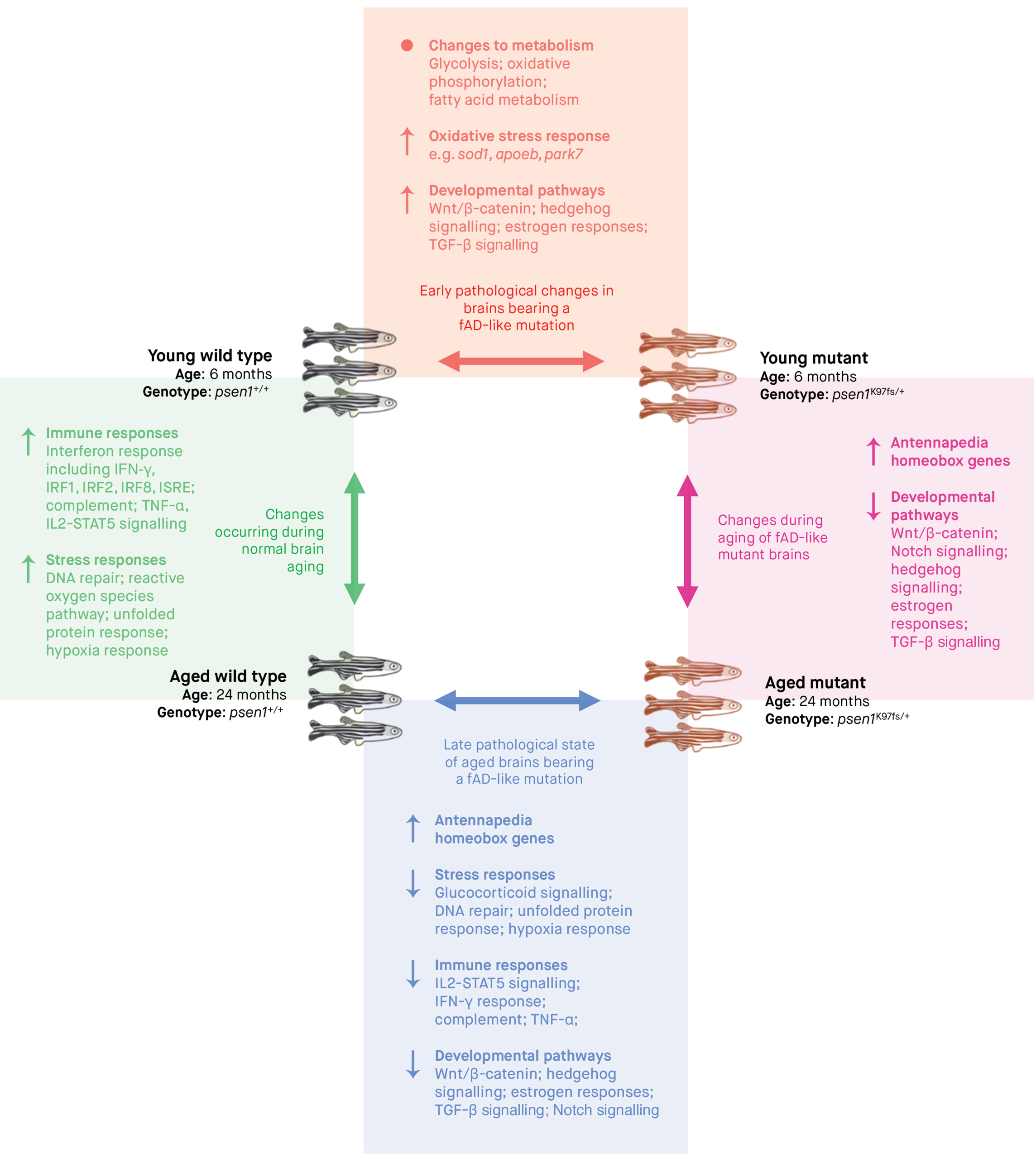
**Summary of the molecular changes in the brains of zebrafish due to aging and/or fAD-like mutation (*psen7*^K97fs/+^)**. For each of the four pairwise comparisons shown, the summarised molecular changes ↑ = overall increased, ↓ = overall decreased, • = significant alterations but not in an overall direction) were inferred from a combination of the following analyses: functional enrichment analysis of differentially expressed genes and proteins, promoter motif enrichment analysis of differentially expressed genes, gene set enrichment analysis of differentially expressed genes, and weighted co-expression network analysis of the gene expression data.

### Evidence of increased stress long preceding AD

We identified a subset of ‘inverted’ genes that are up-regulated in young mutant brains, but down-regulated in aged mutant brains. Although this pattern might be initially overlooked, similar patterns have also been observed in human cases. At the structural-level, the brains of asymptomatic children with *PSEN1* mutations display greater functional connectivity and increased grey matter volumes of several brain regions^20^. Interestingly, adult *PSEN1* mutation carriers who are still asymptomatic retain these increased brain structure sizes, but upon developing AD symptoms, the affected brain structures rapidly decrease in size^50–52^. These changes are likely to be mirrored at the molecular level in the brain. Patients with Mild Cognitive Impairment, pre-clinical AD, or Down Syndrome (who often develop AD in adulthood) initially display increased expression of particular genes, which decreases when AD symptoms become more severe^23^,^53–55^. Collectively, results from these studies and our fAD-like mutant zebrafish suggest that early increases in brain activity likely precede AD symptoms in both *PSEN1*-mutation carriers and more general cases of AD. Evidently, to find strategies for preventing AD progression while patients are still asymptomatic, it is important to understand the causes of this increased brain activity.

Our results suggest that stress responses are likely to contribute to early increases in brain activity for fAD mutation carriers. In mutant zebrafish, the inverted gene expression pattern seems to arise partially from altered glucocorticoid signalling. In humans, chronically increased glucocorticoid signalling in the brain can lead to glucocorticoid resistance, whereby the brain is unable to increase glucocorticoid signalling even during stressful conditions^56^,^57^. We did not confirm whether glucocorticoid signalling and cortisol levels were altered in zebrafish brains *in vivo.* However, many of the inverted genes possess glucocorticoid receptor elements in their promoters, and one particular inverted gene *fkbp5)* encodes a protein which is known to bind directly to the glucocorticoid receptor to negatively regulate its activity. Previous studies in humans demonstrate that *fkbp5* levels are highly responsive to chronic stress and stress-related diseases (e.g. bipolar disorder; depression in AD^58^), implying *that fkbp5* expression is a sensitive marker of glucocorticoid signalling. Our analysis supports this, with *fkbp5* mRNAs showing a significant difference in expression between mutant and wild type brains (logFC = 2.1, FDR-adjusted *p*-value = 1.77e-06 in young mutant vs wild type; logFC = −3.9, FDR-adjusted *p*-value = 3.16e-08 in aged mutant vs wild type). Aside from altered glucocorticoid signalling, we also linked altered gene expression to a range of biological processes that are altered in mutant zebrafish brains. If we assume that these mutant zebrafish accurately model the genetics underlying human AD, then these alterations may offer insight into early changes in the brains of human fAD-mutation carriers and, potentially, other individuals predisposed to AD. The brains of young mutant zebrafish exhibit changes to developmental signalling pathways (Wnt/β-catenin signalling, hedgehog signalling, TGF-β signalling), stress and immune responses (DNA repair, IL2-STAT5 signalling, complement system, IFN-γ response, inflammatory response), hormonal changes (early and late estrogen responses, androgen response), and energy metabolism (glycolysis, oxidative phosphorylation). Evidently, appropriate regulation of these biological processes is critical for brain function, so it is unsurprising that disruption of these processes in the brain has been linked previously to various pathological states, including early stages of neurodegeneration^59–62^. It is difficult to infer how a single heterozygous fAD mutation could alter such diverse processes in the brain. It is unsurprising that cellular processes and pathways where Presenilin1 proteins are directly involved (e.g. Wnt/β-catenin signalling, Notch signalling^63–68^) are altered in the fAD-like mutant zebrafish brains. However, the mechanisms directly linking the fAD-like mutation to other processes (e.g. immune and stress responses) cannot be inferred directly from our results. In future, it would be useful to include zebrafish younger than 6 months of age and additional age groups between 6 and 24 months to further elucidate the sequential progression of brain transcriptome changes in zebrafish carrying a fAD-like mutation.

Quantifying protein abundance in young mutant zebrafish brains revealed additional sources of early-life stress. In the young mutant zebrafish brains, proteins associated with oxidative stress responses and energy metabolism in mitochondria already displayed altered abundance. Overall, stress responses were increased, consistent with the RNA-seq data, and decreased abundance of metabolic and antioxidant proteins indicated potential impairment to mitochondrial function. Both increased oxidative stress and altered energy metabolism appear to be early events in AD^2,69-75^ and it is possible that these events may induce early stress responses in the brain. Unfortunately, although we intended to integrate transcriptome and proteome information more thoroughly in this study, we were only able to reliably quantify 323 proteins across all samples, with most of these being biased towards higher-abundance proteins in the zebrafish brain. Using more sophisticated mass spectrometry technology should provide higher-resolution detection of lower abundance proteins and metabolites^76^.

### Comparison of our elucidation of brain co-expression networks with similar studies

We find that several gene modules are altered in mutant zebrafish brains and in human early-onset AD brains. The most significantly preserved module in the zebrafish and human brain co-expression networks analysed is enriched in immune responses and genes expressed by type 1 and 2 microglia. Remarkably, two independent studies involving co-expression analysis of AD brains by Miller et al. ^44^ and Zhang et al. ^45^ also found a prominent immune-microglia module.

By comparing the zebrafish and human brain co-expression networks, we found several modules of genes which were altered in both mutant zebrafish brains and early-onset AD human brains. We found that the most significantly preserved module across the zebrafish and human brain co-expression networks was enriched in immune responses and genes expressed by type 1 and 2 microglia. Miller et al. ^44^ used 18 human datasets and 20 mouse datasets representing various brain regions, diseases and Affymetrix platforms to construct consensus co-expression networks (aggregated from multiple datasets) for the human and mouse brain. Similarly, the study by Zhang et al. ^45^ used over 1,647 post-mortem brain tissue samples from patients with late-onset AD to construct a consensus co-expression network for the human brain. Compared to our analysis, the large-scale analyses by Miller et al. ^44^ and Zhang et al. ^45^ used many more samples, had broader representation of brain tissue types, included more general cases of sporadic and late-onset AD, and constructed consensus networks to increase the robustness of the co-expression network. Nevertheless, both studies independently identified a module associated with AD which is significantly enriched in immune processes and microglial genes. Collectively, the results from these studies and our analysis here suggest that immune responses and microglial cells are strongly implicated in both early-onset fAD and sporadic late-onset forms of AD. This is reasonable, given that microglia are essential cells in the brain's innate immune response, and neuroinflammation is a prominent feature of AD. Our results also support the idea that these responses are highly conserved across species.

Our analysis reveals additional key insights that may help explain why the immune-microglia module is involved in AD. First, promoter enrichment analysis of genes in the immune-microglia module indicates statistically significant enrichment in several known motifs. Intriguingly, all of these motifs are binding sites for transcription factors from either the ETS (SpiB, ELF3, ELF5, PU. 1, EHF) or IRF (IRF3, IRF8, IRF1) families. This finding is important, because 1) ETS and IRF transcription factor motifs are also enriched in the promoters of genes that are up-regulated with brain aging in wild type zebrafish, but not in genes that are up-regulated with brain aging in mutant zebrafish. This suggests that the genes they regulate are important during normal brain aging and that their dysregulation may contribute to pathology. 2) ETS and IRF transcription factors are known to mediate critical biological functions. The ETS family regulates cellular differentiation, proliferation, cell-cycle control, apoptosis, migration and mesenchymal-epithelial interactions^77^,^78^, while the IRF family regulates immune and other stress responses. Our results are consistent with those in a previous study by Gjoneska et al. ^79^ that analysed RNA-seq and ChIP-seq (chromatin immunoprecipitation sequencing) data from mouse and human brain tissues, which found that immune response genes were up-regulated in both the CK-p25 mouse model and in human sporadic AD, that these genes were enriched in ChIP-seq peaks corresponding to ETS and IRF transcription factor motifs, and that microglia-specific activation was likely responsible for these gene expression changes.

Although revealing valuable insights, our comparison of zebrafish and human brain co-expression networks is limited for several reasons. First, the zebrafish dataset is RNA-seq data while the human dataset is microarray-based data. Differences in the correlation in noise inherent to either technology have been shown previously to affect connectivity within network modules, although functional enrichment of modules tends to be highly preserved between microarray and RNA-seq co-expression networks^39^. This means that our current analysis is likely unable to accurately detect most highly-connected genes within modules that are preserved between the zebrafish and human co-expression networks, which makes it difficult to find key genes that might drive changes in the co-expression network. Second, while entire zebrafish brains were used in our analyses, the analysed human brain tissues were derived only from the posterior cingulate region. Although a previous analysis identified similar co-expression networks across different brain regions^43^, it is likely that differences in the properties of networks constructed from different regions could confound our ability to fully detect similarities and differences between the co-expression networks of zebrafish and human brains.

### AD-like transcriptome and proteome changes can occur without amyloid plaques typically associated with AD

Somewhat surprisingly, the gene and protein expression changes observed in our aged fAD-like mutation model zebrafish were not reflected in an obvious histopathology. However, this is consistent with an attempt to model neuronal ceroid lipofuscinosis in adult zebrafish^47^ and with observations from heterozygous fAD mutation knock-in models in mice^25–27^ (although in general, mouse single heterozygous mutation brain histology phenotypes have not been reported). It is important to realise that differences in scale between the mass of a human brain and the brains of mice and zebrafish, (∼1,000-fold and ∼70,000 fold respectively) mean that any metabolic or other stresses in the small brains of the genetic models are likely exacerbated in the huge human brain^80^. Human brains also lack the regenerative ability of zebrafish, while mice and zebrafish both show sequence divergences in the Aβ regions of their APP orthologous genes greater than seen in most mammals^33^,^81^,^82^. Nevertheless, the heterozygous fAD-like mutation models of mice and (with this paper) zebrafish are probably the closest one can come to modelling AD in these organisms without subjectively imposing an opinion of what AD is by addition of further mutations, transgenes etc. We speculate that the transcriptional state of 2-year-old fAD-like mutant zebrafish brains may represent, at the molecular level, the zebrafish equivalent of AD.

It is important to remember that the pathological role in AD of Aβ, neuritic plaques, and neurofibrillary tangles is still debated and that around one quarter of people clinically diagnosed with AD are, upon post-mortem examination, seen to lack typical amyloid pathology^83^. By the current definition, these people do not have AD^84^ although this restrictive definition has been questioned^85^. Many people also have brains containing high levels of Aβ^86^ or Braak stage III to VI neurodegeneration^83^ without obvious dementia. Thus, the connection between amyloid pathology, histopathological neurodegeneration and Alzheimer’s disease dementia is unclear. Our data indicate that the AD cellular pathologies may occur subsequent to cryptic but dramatic changes in the brain’s molecular state (gene and protein expression) that are the underlying drivers of AD.

Finally, it is also important to acknowledge that the specific fAD mutation (K115fs) modelled in this study is an uncommon fAD mutation that causes frameshifting and truncation of the human PSEN2 protein, and it is unclear how gene expression alterations caused by this mutation may differ from the far more common frame-preserving fAD mutations. Our laboratory is currently developing additional mutant zebrafish modelling different fAD-causing mutations, and future analysis incorporating these zebrafish to produce a consensus co-expression network should help to identify and refine a “signature” of the gene and protein expression changes that cause fAD.

## Conclusion

Overall, our study highlights the importance of studying gene expression changes in animals closely modelling the genetic state of human fAD in order to elucidate the early molecular-level brain changes driving AD pathogenesis. In particular, zebrafish heterozygous for single, endogenous fAD-like mutations may be useful for exploring several AD-related changes in the brain, including increased brain activity preceding AD symptoms, the role of glucocorticoid-mediated stress responses in the development of AD, and the roles of ETS and IRF transcription factors in regulating microglia-associated genes in AD.

## Methods

### Zebrafish husbandry and animal ethics

Tubegin zebrafish were maintained in a recirculated water system. All work with zebrafish was conducted under the auspices of the Animal Ethics Committee and the Institutional Biosafety Committee of the University of Adelaide.

### Generation of TALEN coding sequences and single stranded oligonucleotide

TALEN coding sequences were designed by, and purchased from, Zgenebio. The DNA binding sites for the TALEN pair targeting *psen1* were (5’ to 3’): left site, CAAATCTGTCAGCTTCT and right site, CCTCACAGCTGCTGTC (Figure 1A_3_ in Supplementary Data). The coding sequences of the TALENs were provided in the pZGB2 vector for mRNA *in-vitro* synthesis. The single stranded oligonucleotide (ssoligo) sequence was designed such that the dinucleotide ‘GA’ deletion was in the centre of the sequence with 26 and 27 nucleotides of homology on either side of this site (Figure 1A_3_). The ssoligo was synthesized by Sigma-Aldrich and HPLC purified. The oligo sequence was (5’ to 3’): CCATCAAATCTGTCAGCTTCTACACACAAGGACGGACAGCAGCTGTGAGGAGC (Figure 1A).

### In-vitro mRNA synthesis

Each TALEN plasmid was linearized with *Not* I. Purified linearized DNA was used as a template for *in-vitro* mRNA synthesis using the mMESSAGE mMACHINE SP6 transcription kit (Thermo Fisher, Waltham, USA) as per the manufacturer’s instructions as previously described^87^.

### Microinjection of zebrafish embryos

Embryos were collected from natural mating and at the 1-cell stage were microinjected with a ∼3nl mixture of 250ng/µl of left and right TALEN mRNA and 200ng/µl of the ssoligo.

### Genomic DNA extraction of zebrafish tissue

#### Embryos

A selection of 10-20 embryos were collected at 24hpf and placed in 150µl of a 50mM NaOH 1×TE solution and then incubated at 95°C until noticeably dissolved (10-20mins). The lysis solution was cooled to 4°C and 50µl of Tris solution (pH 8) was added. The mixture was then centrifuged at maximum speed for 2 mins to pellet cellular debris. The supernatant was transferred into a fresh microfuge tube ready for subsequent PCR.

#### Adult fin clips

For fin clips, adult fish were first anesthetised in a 0.16mg/mL tricaine solution and a small section of the caudal fin was removed with a sharp blade. Fin clips were placed in 50µl of a 1.7µg/ml Proteinase K 1×TE solution and then incubated at 55°C until noticeably dissolved (2-3hours). The lysis solution was then placed at 95°C for 5mins to inactivate the Proteinase K.

## Genomic DNA PCR and sequencing for mutation detection

To genotype by PCR amplification, 5µl of the genomic DNA was used with the following primer pairs as relevant. Primers to detect wild type (WT) sequence at the mutation site: primer psen1WTF: (5’TCTGTCAGCTTCTACACACAGAGG3’) (GA nucleotides in italics) with primer psen1WTR: (5’AGTAGGAGCAGTTTAGGGATGG3’). Primers to detect the presence of the GA dinucleotide deletion: primer psen1GAdelF: (5’AATCTGTCAGCTTCTACACACAAGG3’) with primer psen1WTR. To confirm the presence of the GA dinucleotide deletion mutation by sequencing of extracted genomic DNA, PCR primers were designed to amplify a 488bp region around the GA mutation site: primer psen1GAsiteF: (5’GGCACACAAGCAGCACCG3’) with primer psen1GAsiteR: (5’TCCTTTCCTGTCATTCAGACCTGCGA3’). This amplified fragment was purified and sequenced using the primer psen1seqF: (5’ AGCCGTAATGAGGTGGAGC 3’). All primers were synthesized by Sigma-Aldrich. PCRs were performed using GoTaq polymerase (Promega, Madison, USA) for 30 cycles with an annealing temperature of 65°C (for the mutation-detecting PCR) or 61°C (for the WT sequence-detecting PCR) for 30 s, an extension temperature of 72°C for 30 s and a denaturation temperature of 95°C for 30 s. PCR products were assessed on 1% TAE agarose gels run at 90V for 30mins and subsequently visualized under UV light.

## Whole brain removal from adult zebrafish

Adult fish were euthanized by placement in an ice water slurry for ∼30 seconds. The whole brain was removed and either RNA or protein was extracted immediately. All fish brains were sampled at late morning/noon to avoid effects of circadian rhythms.

## RNA extraction from whole brain

Total RNA was isolated from mutant and WT siblings using the *mir*Vana miRNA isolation kit (Thermo Fisher). RNA isolation was performed according to the manufacturer’s protocol. First the brain is lysed in a denaturing lysis solution. The lysate is then extracted once with acid-phenol:chloroform leaving a semi-pure RNA sample. The sample is then purified further over a glass-fiber filter to yield the total RNA. This procedure has been formulated specifically for miRNA retention to avoid the loss of small RNAs. Total RNA was then sent to the ACRF Cancer Genomics Facility (Adelaide, Australia) to assess RNA quality and for subsequent RNA sequencing.

## Protein extraction and proteomic analysis of adult brain

### Sample preparation

Freshly removed adult zebrafish brains were lysed under denaturing conditions in 7 M urea (Merck) plus complete protease inhibitors (Roche) using a Bioruptor (Diagenode, Seraing, Belgium) in ice cold water. Samples were quantified using the EZQ protein assay (Life Technologies) and the extracts were trypsin-digested using the FASP method^88^. Protein samples were then sent to the Adelaide Proteomics Centre (Adelaide, Australia) for quantification and data acquisition.

### Data Acquisition

Nano-LC-ESI-MS/MS was performed using an Ultimate 3000 RSLC system (Thermo Fisher Scientific) coupled to an Impact HD™ QTOF mass spectrometer (Bruker Daltonics, Bremen, Germany) via an Advance Captive Spray source (Bruker Daltonics). Peptide samples were pre-concentrated onto a C18 trapping column (THC164535, Thermo Fisher) at a flow rate of 5 µL/min in 2% (v/v) ACN 0.1% (v/v) FA for 10 minutes. Peptide separation was performed using a 75µm ID 50 cm C18 column (THC164540, Thermo Fisher) at a flow rate of 0.2 µL/minutes using a linear gradient from 5 to 45% B (A: 5% (v/v) ACN 0.1% (v/v) FA, B: 80% (v/v) ACN 0.1% (v/v) FA) over 180 minutes. MS scans were acquired in the mass range of 300 to 2,200 m/z in a data-dependent fashion using Bruker’s Shotgun Instant Expertise™ method (singly charged precursor ions excluded from acquisition, CID from 23% to 65% as determined by the m/z of the precursor ion).

### Data Analysis

The acquired peptide spectra were identified and quantified using the mass spectrometry software MaxQuant with the Andromeda search engine against all entries in the non-redundant UniProt database (protein and peptide false discovery rate set to 1%). The MaxQuant software allows for the accurate and robust proteomewide quantification of label-free mass spectrometry data^89^.

## RNA-seq Analysis

### Data Processing

We used *FastQC^90^* to evaluate the quality of the raw paired-end reads and identified several issues with read quality, adapter sequences, GC content, and over-represented sequences. Using *AdapterRemoval^91^,* we trimmed, quality-filtered and removed adapter sequences from paired-end reads of each RNA-seq library, resolving issues related to read quality and adapter sequences. From the *FastQC* reports, we realised that some over-represented sequences in the raw and trimmed reads corresponded to ribosomal RNA, possibly from insufficient depletion during RNA-seq library preparation. We removed ribosomal RNA sequences *in silico* by aligning all trimmed reads to known zebrafish ribosomal RNA sequences followed by discarding all reads that aligned. Next, we used *HISAT2^92^* to align reads to the reference zebrafish genome assembly (GRC_z_10) downloaded from Ensembl. Using *Picard*^93^ and the MarkDuplicates function, we removed optical and PCR duplicates from the aligned reads. Following de-duplication, *FastQC* analysis revealed that de-duplication resolved the issues relating to GC content and over-represented sequences. Finally, to quantify gene expression, we used *FeatureCounts* to count the reads aligning to each Ensembl gene model^94^. The output of *FeatureCounts* gave a matrix of gene expression counts for 32,266 genes for each of the 12 RNA-seq libraries.

### Differential Gene Expression Analysis

Differential gene analysis was conducted in R^95^ using the packages *edgeR*^96^ and *limma*^97–99^. We filtered out genes expressed at insufficient levels to be informative, retaining genes with >1.5 counts per million in at least 6 of the 12 RNA-seq libraries. This reduced the number of genes in the analysis from 32,266 to 18,296. We then calculated TMM-normalisation factors to account for differences in library sizes, and applied the RUVs method from the *RUVseq* package^100^ to account for a batch effect with one factor of unwanted variation (*k* = 1). Differential gene expression analysis was performed using *limma.* We considered genes differentially expressed if the FDR-adjusted *p*-value associated with their moderated *t*-test was below 0.05. We used the *pheatmap* R package^101^ to produce all heatmaps.

### Gene Set Testing

We downloaded the Hallmark gene set collection from the Molecular Signature Database (v6.1)^36^ as a .gmt file containing genes with human Entrezgene identifiers. Using *biomaRt^102^,^103^,* we converted human Entrezgene identifiers to zebrafish Entrezgene identifiers. To perform gene set testing, we applied the fast rotation gene set testing (FRY) method^104^ for each comparison in the contrasts matrix. We considered all gene sets for a particular comparison with Mixed FDR < 0.05 as differentially expressed. To obtain estimates of the proportions of up-regulated and down-regulated genes for each significant gene set, we used the ROAST^105^ method with 9,999 rotations and set the ‘set.statistic’ option set to ‘mean’ to maintain consistency with the results obtained from FRY. We performed FDR multiple testing adjustment for the results from FRY and ROAST.

### Promoter Motif Analysis

We performed promoter motif enrichment analysis using *HOMER^106^,^107^.* We downloaded a set of 364 zebrafish promoter motifs collated by *HOMER* authors from published ChIP-seq experiments using the command `configureHomer.p1-install zebrafish-p`. We retained default parameters with the *HOMER* `findMotifs.pl` program with the following modifications: the 18,296 ENSEMBL genes used in the differential gene expression analysis were specified as the background genes, and promoter regions were defined as 1500 bp upstream and 200 bp downstream of the transcription start site of each gene. We defined motifs as being significantly enriched in a set of genes if the Bonferroni-adjusted *p*-value was less than 0.05.

## Proteomic Analysis

### LC-MS/MS Data Processing

Raw MS/MS spectra were analysed using *MaxQuant* (V. 1.5.3.17). A False Discovery Rate (FDR) of 0.01 for peptides and a minimum peptide length of 7 amino acids was specified. MS/MS spectra were searched against the zebrafish UniProt database. *MaxQuant* output files for the 6-month-old and 24-month-old samples were processed in separate batches with the *MSStats* R package^108^ due to an unresolvable batch effect in the data caused by generation of spectra on different days by different operators. Briefly, peptide intensities were log**2**-transformed and quantile normalised, followed by using an accelerated failure time model to impute censored peptides. Peptide-level intensities were summarised to protein-level intensities using a Tukey's median polish method. This resulted in 2,814 peptides (summarised to 534 proteins) for the 6-month-old data and 3,378 peptides (summarised to 582 proteins) for the 24-month-old data. After *MSStats* data processing, the 6-month-old and 24-month-old protein log_2_-intensities were combined, quantile normalised using the `normaliseBetweenArrays` function from the *limma* R package, and filtered to only retain the 323 proteins that were detected across all samples.

### Differential Protein Analysis

Differential protein abundance analysis was performed using *limma* which has been shown to be highly applicable to proteomics data^109^. A linear model was fitted to each protein and moderated t-test performed to test for differential abundance between samples. Proteins were identified as being differentially abundant if their FDR-adjusted *p*-values were below 0.05. Over-representation analysis using the `goana` and `kegga` functions from *limma* were used to test for enriched gene ontology terms and KEGG pathways respectively.

## Network Analysis

### Network construction

Network construction and analysis used functions from the *WGCNA* R package^40^ applied to the same filtered set of gene counts from the earlier differential expression analysis. We additionally downloaded processed human microarray data from ArrayExpress under the E-GEOD-39420 accession number (raw dataset at GEO with accession number GSE39420). This data set contains gene expression profiles derived from the posterior cingulate tissue of 21 adult brains (7 *PSEN1* mutation carriers with early-onset AD; 7 early-onset AD cases without *PSEN1* mutations; 7 controls). Mutations represented in the *PSEN1* mutation carriers include E120G, V89L, and M139T. The mean age was 53.9 years (s.d = 6.5 years) for the *PSEN1* mutation carriers with early-onset AD, 63.3 years (s.d = 4.3 years) for the early-onset AD lacking *PSEN1* mutations, and 49.7 years (s.d = 4.5 years) for the control group. Principal component analysis revealed no association between having AD and either sex or *APOE* genotype of individuals. The microarray data was already background adjusted, normalised and summarised by Antonell et al ^24^. We then applied the collapseRows function with default parameters to retain only one gene per probe^110^. Using the *BioMart* package, we retrieved zebrafish ‘one-to-one’ homologous genes for human genes. We matched zebrafish genes to human homologous genes via common Ensembl gene identifiers and retained only genes that were expressed in both the human and zebrafish datasets, leaving 13,110 genes for network construction. To further reduce noise during network construction, we calculated connectivities for each gene in each dataset and retained only the 7,118 genes with connectivities above the 10th percentile of all connectivities. To construct approximately scale-free weighted networks, the Pearson correlation was calculated between each pair of genes, and the resulting correlation matrix was raised to the soft-thresholding power of 14 to produce a signed adjacency matrix for each dataset^40^. Next, we applied a transformation to obtain a measure of topological overlap for each pair of genes. Lastly, we hierarchically clustered genes in each dataset based on their topological overlap (using the measure 1 - Topological Overlap) to form a dendrogram where branches represented distinct modules of genes showing similar expression patterns across all samples. To identify modules of co-expressed genes, we used the Hybrid Tree Cut method from the *dynamicTreeCut* package^111^. We used default parameters except for the following modifications to define modules in the zebrafish network: minimum module size set at 40 genes, 0.90 as the maximum distance to assign previously unassigned genes to modules during PAM (Partioning Around Medoids) stage, and the deepSplit parameter to 1 for the zebrafish data set and 2 for the human data set.

### Network Analysis

We assessed the functional enrichment of each module using default settings in the *anRichment* R package. To calculate the correlation between modules and phenotypic traits, we calculated the hybrid-robust correlation between the first principal component of each module and four binary variables defining the experimental conditions^112^. We evaluated the preservation of zebrafish modules in the human network and vice versa using the “modulePreservation” function from *WGCNA,* which uses a permutation-based approach to determine whether module properties (e.g. density, connectivity) are preserved in another network^46^. We also used the Sankey diagram functionality in the *networkD3* package to visualise overlap between zebrafish and human modules^113^.

### Network visualisation

To visualise zebrafish and human networks, we imported edges and nodes into *Gephi* and applied the OpenOrd algorithm with default settings, which is suitable for visualising weighted, undirected networks^114^. We coloured the nodes (genes) based on their assigned modules from *WGCNA.*

### Code availability

Source code of all R analyses and associated data is available at github.com/UofABioinformaticsHub/k97fsZebrafishAnalysis.

### Data availability

Raw RNA-seq data is available on the European Nucleotide Archive (ENA) with the accession number PRJEB24858.

## Acknowledgements

We thank the Carthew Foundation for financial support of our laboratory, and Awais Choudhry for helping with the promoter motif analysis.

## Author contributions

NH performed the bioinformatics analysis and drafted the manuscript. MN generated the mutant fish, isolated brain RNA and protein, drafted the description of this and edited the manuscript. AD performed histological analysis of 24-monthold brains under JK’s supervision. JK drafted the description of this. AL performed initial promoter analysis. X-FZ and NM performed initial histological analysis of 6- month-old brains. AL wrote and guided the RNA-seq data processing pipeline and contributed ideas for improving RNA-seq data quality. DA contributed bioinformatics supervision and ideas for the analysis. SP directly supervised NH, guided the analysis and performed quality-control checks on results and edited the manuscript, ML conceived the project and supervised all aspects of it and edited the manuscript. All co-authors critically read and approved the manuscript.

